# LOX-1^+^ immature neutrophils predict severe COVID-19 patients at risk of thrombotic complications

**DOI:** 10.1101/2020.09.15.293100

**Authors:** Béhazine Combadiere, Lucille Adam, Paul Quentric, Pierre Rosenbaum, Karim Dorgham, Olivia Bonduelle, Christophe Parizot, Delphine Sauce, Julien Mayaux, Charles-Edouard Luyt, Alexandre Boissonnas, Zahir Amoura, Valérie Pourcher, Makoto Miyara, Guy Gorochov, Amélie Guihot, Christophe Combadière

## Abstract

**Rational:** Lymphopenia and neutrophil/lymphocyte ratio may have prognostic value in coronavirus disease 2019 (COVID-19) severity.

**Objective:** We sought to investigate the representation of neutrophil subsets in severe and critical COVID-19 patients based on Intensive Care Units (ICU) and non-ICU admission.

**Methods:** We developed a multi-parametric neutrophil profiling strategy based on known neutrophil markers to distinguish COVID-19 phenotypes in critical and severe patients.

**Results:** Our results showed that 80% of ICU patients develop strong myelemia with CD10^−^CD64^+^ immature neutrophils. Cellular profiling revealed two distinct neutrophil subsets expressing either the lectin-like oxidized low-density lipoprotein receptor-1 (LOX-1) or the Interleukin-3 receptor alpha (CD123), both significantly overrepresented in ICU patients compared to non-ICU patients. The proportion of LOX-1-expressing immature neutrophils positively correlated with clinical severity, with the cytokine storm (IL-1*β*, IL-6, IL-8, TNF*α*), and with intravascular coagulation. Importantly, high proportions of LOX-1^+^-immature neutrophils are associated with high risks of severe thrombosis.

**Conclusions:** Together these data suggest that point of care enumeration of LOX-1-immature neutrophils might help distinguish patients at risk of thrombosis complication and most likely to benefit from intensified anticoagulant therapy.

## Introduction

Since the first reports of an outbreak of a severe acute respiratory syndrome caused by coronavirus 2 (SARS-CoV-2) in China in December 2019 (1, 2), the coronavirus disease 2019 (COVID-19) has grown to be a global public health emergency, with cases of COVID-19 around the world reaching 8 385 440 cases and 450 686 deaths as of June 19^th^, 2020 (for up-to-date, https://www.who.int/emergencies/diseases/novel-coronavirus-2019). SARS-CoV-2 infection is characterized by a range of symptoms including fever, cough, fatigue and myalgia in the majority of cases and occasional headache and diarrhea (1, 3). Among reported cases, approximatively 80% present with mild condition, 13% serious condition, and 6% develop a critical state requiring intensive care, the latter associated with a fatality rate of 2 to 8% of reported cases (4). Some severe cases of COVID-19 progress to acute respiratory distress syndrome (ARDS), accountable for high mortality related to the damages of the alveolar lumen. Numerous patients with ARDS secondary to COVID-19 develop life-threatening thrombotic complications (5).

Previous coronaviruses-related infections have been characterized by the onset of a cytokine storm (6). It is therefore reasonable to postulate that the inflammatory response measured both at cellular and molecular levels could represent a strong prognostic signature of the disease. Molecular assays have been the gold standard to directly detect the presence of the virus as well as to characterize the infection onset. The cytokine storm remains as of today an uncontrollable inflammatory response leading to viral sepsis, acute respiratory distress syndrome, respiratory failure, shock, organ failure or death (7, 8). In addition to the life-threatening course of the disease, many critically ill patients develop complications with a high burden (e.g. thrombosis, ventilator-associated pneumonia, acute kidney injury…). Strong predictive markers are still missing for these.

A retrospective cohort of 201 patients with confirmed COVID-19 pneumonia revealed that older age, neutrophilia, organ and coagulation dysfunction were the major risk factors associated with the development of ARDS and progression to death (9). ARDS and sepsis are frequent complications among deceased patients (10). In severe cases, bilateral lung involvement with ground-glass opacities is the most common chest computed tomography (CT) finding. More surprisingly, abnormal CT scans were also reported on asymptomatic COVID-19 patients (11). Immune transcriptome profiling of bronchoalveolar lavage fluids of COVID-19 patients also displayed high levels of pro-inflammatory cytokines (6). In addition, serum concentrations of both pro- and anti-inflammatory cytokines including IL-6, TNFα, and IL-10, were increased in the majority of severe cases and were markedly higher than those of moderate cases, suggesting that the cytokine storm might be associated with disease severity and leading the way to the development of potential immune-modulatory treatments (3, 12). The cytokine storm is associated with a massive influx of innate immune cells, namely neutrophils and monocytes, which could worsen lung injury. However, little is known about the innate immune features and the molecular mechanisms involved in COVID-19 severity.

Increasing clinical data indicated that the neutrophil-to-lymphocyte ratio (NLR) was a powerful predictive and prognostic indicator of severe COVID-19 (13–15). Lymphopenia, neutrophilia, and high NLR are associated with a more severe viral infection (13, 16).

We previously identified two new CD10^−^CD64^+^ neutrophil subsets expressing either PD-L1 or CD123 that were specific to bacterial sepsis (17). In addition to these markers, previous work showed that LOX-1 is an important mediator of inflammation and neutrophils dysfunction in sepsis and cancers (18, 19). To test the hypothesis of a virally-driven neutrophil profile, we developed a multi-parametric neutrophil profiling strategy based on known neutrophil markers to distinguish COVID-19 phenotypes in critical and severe patients.

## Materials and Methods

### Study participants

Fresh blood samples from 38 consecutive adult patients with COVID-19 referred to the Department of Internal Medicine 2, Department of Infectious Diseases and Intensive Care Units (ICU), Pitié-Salpêtrière Hospital, Paris were included in the study between March 18, 2020 and April 29, 2020. The diagnosis of COVID-19 relied on SARS-CoV-2 carriage in the nasopharyngeal swab, as confirmed by real-time reverse transcription-PCR analysis. Demographic and clinical characteristics are detailed in Supplementary Table S1. All flow cytometric analyses were performed on fresh whole blood cells collected at the admission to the hospital and ICU. Sera were collected for cytokine measurement.

### Study approval

The study was conducted in accordance with the Declaration of Helsinki and the International Conference on Harmonization Good Clinical Practice guidelines and approved by the relevant regulatory and independent ethics committees. All patients gave oral informed consent. The study was registered and approved by local ethical committee of Sorbonne-Université/assistance-publique hopitaux de Paris for standard hospitalized patients (N°2020-CER2020-21) and ICU patients (N° CER-2020-31).

### Flow cytometry

One hundred µl of fresh whole blood collected on Anticoagulant Citrate-Dextrose solution (ACD) for patients in intensive care or on EDTA for patients in standard hospitalization were stained with a mix of monoclonal antibodies. Samples were diluted in brilliant violet buffer (BD biosciences) and incubated 20 min at room temperature in the dark. The antibody panel (Supplementary Table S2) included: CD15-BV786, CD14-BUV737, CD10-BUV395 (BD, Le Pont de Claix, France); CRTH2-FITC, CD123-PE, LOX-1-BV421, CD64-BV605 and PD-L1-BV711 (Biolegend, San Diego, USA). One ml of BD FACS lysing (BD biosciences) solution 1X was directly added to the cells to lyse red blood cells, incubated 20min, centrifuged, and washed with PBS. Leukocytes were resuspended in PBS before analysis with a BD LSR FORTESSA X-20. FlowJo software 10.0 was used for analysis of marker expression on neutrophils. One hundred µl of whole blood were additionally stained for several patients with an FMO mix missing of antibodies targeting CD123, LOX-1, and PD-L1, in order to determine the threshold of expression of these markers.

### Data presentation and statistical analysis

Statistical analyses of the immunological data and graphic representations were performed with Prism 8.0 (GraphPad Software Inc.). Two-tailed Student’s t-test were used for group comparisons, and one-way and two-way ANOVA tests with Bonferoni for multiple comparison tests. The potential association between serum cytokine levels or marker-expressing neutrophils frequencies was evaluated using Spearman correlation (one-tailed), with significance defined by a p-value < 0.05: * for p < 0.05; ** for p < 0.01; *** for p < 0.001; **** for p < 0.0001. Principal Component Analyses were performed with R software 3.3.1.

## Results

### Demographics and baseline characteristics of ICU and non-ICU COVID-19 patients

Thirty-eight COVID-19 patients admitted to either ICU departments or non-ICU departments were included. SARS-CoV-2 infection was confirmed on nasopharyngeal swab by positive RT-PCR in accordance with WHO interim guidance. Clinical and biological characteristics of the 38 patients are shown in Supplementary Table S1. Median age was 57 years (range 25-79 years), with 65.8% of them being males. Analysis was performed on median average 8 days after the onset of symptoms (median was 8 days for the ICU patients, 13 days for the non-ICU patients). The most common past medical comorbidities were hypertension (50%), type 2 diabetes (34.2%) and obesity (36.8%). Treatment regimen at baseline was mostly anti-hypertensive therapy (ACE inhibitors 26.3% and angiotensin II receptor blockers 15.8%). Severity at baseline was assessed by the SAPS II score for all patients (median 33, ranging from 25 to 78) and an additional SOFA score for ICU patients (median 8.5, ranging from 2 to 17). Twenty-eight patients were assessed with CT chest imaging, with ground-glass opacities and/or consolidation > 50% of the lung field among 50% of all patients, with up to 81.3% of the ICU patients. Laboratory findings showed a decreased median lymphocyte count at 0.94×10^9^/L, an increased median neutrophil count at 7.87×10^9^/L, an increased median lactate dehydrogenase at 475.5 U/L and an increased median D-dimer level at 2450 ng/ml. During hospitalization, 8 patients received hydroxychloroquine (42.1%), while all patients received antibiotics. Oxygen therapy was administered to 100% of patients; ICU patients were ventilated with invasive mechanical ventilation for 87.5% of them, while 54.2% received extracorporeal membrane oxygenation. Acute respiratory distress syndrome occurred among 55.3% of all patients (87.5% of ICU patients) and acute kidney injury among 31.2% of all patients. Among the 38 patients, 2 patients were diagnosed with pulmonary embolism (5.3%) and 10 patients (all ICU) (26.3%) were diagnosed with venous thromboembolism. 76.3% of all patients were discharged as of June 8, 2020, while 10.5% remained in hospital and 13.2% had died, the latter all being ICU patients.

### Increased proportions of circulating immature neutrophils expressing either CD123 or LOX-1 in severe COVID-19 patients

In order to identify neutrophil surface markers that may help to predict the severity of the infection at hospital admission, we designed an observational study including 38 individuals and analyzed their neutrophil phenotypes, comparing them among patients admitted to ICU (n=24) or not (n=14) within the first day following their admission (Supplementary Table S1). COVID-19 patients from ICU displayed more severe clinical and biological signs than non-ICU patients, with an elevated Simplified Acute Physiology Score (SAPS II; 35.5, n=24 and 25.5, n=14; p=0.056), higher serum lactate dehydrogenase (504, n=24 and 324, n=14; p=0.005), and higher D-dimers (2760, n=23 and 1860, n=12; p=0.25). However, they did not differ while comparing neutrophils counts (8.75, n=24 and 5.185, n=14; p=0.06) and level of lymphopenia (0.925, n=24 and 1.185, n=14; p=0.41). Patients characteristics confirmed previously published data with notably a high prevalence of obese patients in COVID-19 patients. Main differences between ICU and non-ICU patients reflected case severity with a high proportion of patient with lung lesions (as observed by ground-glass opacities on chest CT) requiring mechanical ventilation and resulting in high in-hospital mortality.

Whole blood immunostaining was performed, within 3h after blood drawing, using a previously published panel designed to give a precise evaluation of immature circulating neutrophils (17). Neutrophils were automatically identified and visualized using Visualization of t-Distributed Stochastic Neighbor Embedding (viSNE implementation of t-SNE) in order to define an imprint for each sample group (Figure 1A). Using an unsupervised classification, neutrophils from ICU patients were organized in the upper left quadrant of the map, whereas the non-ICU patients’ neutrophils were in the upper right quadrant.

**Figure 1:**
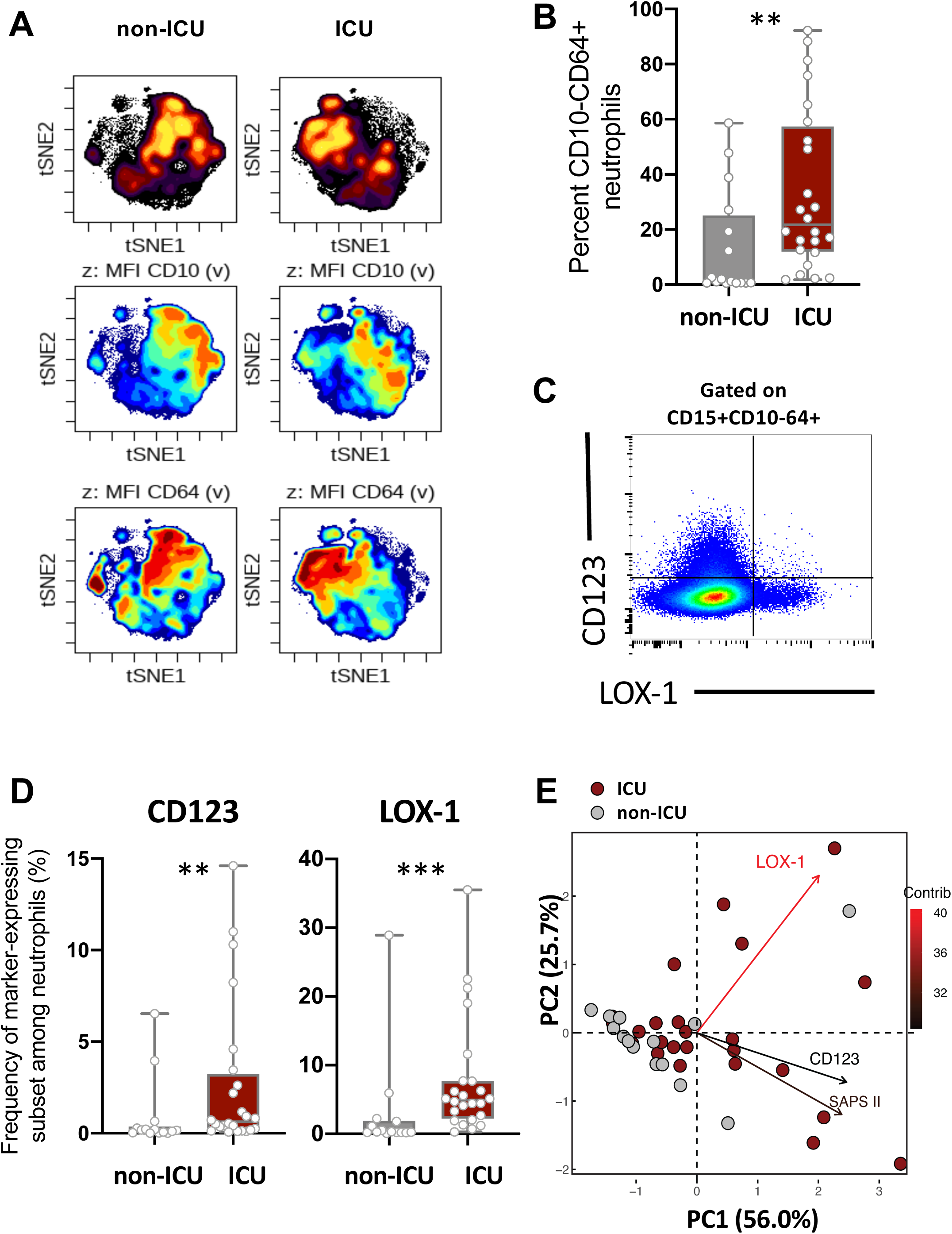
Severe COVID-19 patients displayed increased immature neutrophil subsets expressing CD123 or LOX-1. (A) viSNE analysis was performed on neutrophils from all samples with cells organized along t-SNE-1 and t-SNE-2 according to per-cell expression of CD15, CD10, CD64, LOX-1, CD123 and PD-L1. Cell density for the concatenated file of each patient’s group (ICU vs Non-ICU) is shown on a black to yellow heat scale. Neutrophils’ CD10, CD64 markers expression is presented on a rainbow heat scale in the t-SNE map of each group concatenated file. (B) Box plots representation (min to max distribution) of CD10^−^CD64^+^ neutrophil subset abundancy among total neutrophils of each group samples. (C) Representative expression of LOX-1 and CD123 on CD10^−^CD64^+^ neutrophils. (D) Abundancy of CD10^−^CD64^+^ neutrophil expressing CD123 or LOX-1 in ICU and non-ICU patients’ groups. identify the median and min to max distribution. Nonparametric Mann-Whitney test was used to compare differences in cellular abundance of neutrophil subsets between groups, with significance defined by a p-value < 0.05: * for p < 0.05; ** for p < 0.01; *** for p < 0.001. (E) Principal component analysis (PCA) using LOX-1+, CD123+ CD10^−^CD64^+^ neutrophil abundancy and SAPS II variables on sample sizes: ICU=24 (dark red circles), non-ICU=14 (grey circles). Percent contribution (contrib) of each variable is indicated in color gradient black-red of the arrows.

This analysis allowed precise delimitation of two main subsets of neutrophils based on the expression of CD10 and CD64 markers; the ICU-abundant upper left area composed of neutrophils with mid-to-low expression of CD10 and high expression of CD64, and the non-ICU-abundant upper right area composed of high-to-mid expression of CD10 and high-to-mid expression of CD64. We next determined whether the identified neutrophil signature would be confirmed using conventional analysis applicable by experts. Neutrophils were identified on the CD15 neutrophil marker and excluding prototypical markers of eosinophil and monocyte, respectively CRTH2 and CD14 (Supplementary Figure S1). Expert-gating strategy confirmed the high abundance of CD10^−^ CD64^+^ neutrophils among ICU patients compared non-ICU patients (Figure 1B and Supplementary Figure S2 for cell counts). Then, we compared the expression of CD123, LOX-1 and PD-L1 surface molecules, formerly known as dysregulated in sepsis (17). All three of them were barely co-expressed on neutrophils and lead to the identification of three distinct neutrophil subpopulations (Figure 1C and data not shown). Subsets of immature neutrophils expressing either CD123 or LOX-1 were more abundant in ICU than in non-ICU patients unlike those expressing PD-L1 (Figure 1D and Supplementary Figure S2 for cell counts). Principal component analysis (Figure 1E) revealed that both CD123 and LOX-1 expressing immature neutrophils contributed independently to patients’ severity as appreciated by SAPS II regardless of age, obesity and other potential confounding factors. These data suggested that immature neutrophil subsets expressing CD123 or LOX-1 may contribute to the severity of the disease.

### LOX-1+ immature neutrophils are positively correlated with clinical severity and cytokine levels

COVID-19 patients from ICU were segregated into two groups based on severity at the time of admission. Patients with low SOFA (<8) had significantly fewer CD123- and LOX-1-expressing immature neutrophils than patients with high SOFA (≥8), (p<0.01 and p<0.001 respectively) (Figure 2A). The abundancy of LOX-1-expressing immature neutrophils correlated positively with the SOFA score (Table 1), with inflammatory cytokines such as IL-1β, IL-6, IL-8, TNFα and with the anti-inflammatory cytokine IL-10, whereas it correlated negatively with IFNα and the multipotent hematopoietic growth factor IL-3. In contrast, proportion of CD123-expressing immature neutrophils did not correlate with any of these cytokines but with IL-17, IL-18, IL-22, while negatively correlating with IFNβ (Table 1). In addition, we observed that the expression of LOX-1 significantly correlated with serum D-dimer concentrations. Principal Component Analysis confirmed that the ICU patients’ severity was associated with the proportion of CD123- and LOX-1-expressing immature neutrophils and distinct patterns of cytokines (Figure 2B). These results lead to the identification of three profiles: 1) patients with high LOX-1+ immature neutrophils proportions and high IL-1β, IL-6, IL-8, TNFα serum levels, 2) patients with CD123+ immature neutrophils and IL-18, IL-22, IFNγ secretion, and 3) a bulk of the patients with lower severity associated with high type-1 interferons levels. These data suggested that immature neutrophil subsets expressing either CD123 or LOX-1 may define a specific profile of severity associated with high levels of pro-inflammatory cytokines.

**Table 1:** Correlation analyses between serum cytokine levels and marker-expressing neutrophil abundancies.

| Variables | Frequency of either CD123 or LOX-1-expression among neutrophils |  |  |  |
| --- | --- | --- | --- | --- |
|  | CD123 |  | LOX-1 |  |
|  | *r* | *P-value* | *r* | *P-value* |
| SOFA | 0.69 | <0.0001**** | 0.68 | 0.0001*** |
| D-dimers |  | ns | 0.42 | 0.023* |
| IL-1β | | ns | 0.56 | 0.003** |
| IL-6 |  | ns | 0.48 | 0.010* |
| IL-8 |  | ns | 0.48 | 0.009 |
| TNFα | | ns | 0.42 | 0.022* |
| IL-10 |  | ns | 0.43 | 0.020* |
| IL-17 | 0.06 | 0.035* |  | ns |
| IL-18 | 0.22 | 0.009** |  | ns |
| IL-22 | 0.32 | 0.005** |  | ns |
| IFNα | | ns | | ns |
| IFNβ | -0.45 | 0.035* | -0.45 | 0.023* |
| IFNγ | | ns | | ns |
| GM-CSF |  | ns |  | ns |
| IL-3 |  | ns | -0.44 | 0.020* |

The Simoa<sup>TM</sup> (single molecule array) HD-1 analyser was used for ultrasensitive multiplex immunodetection of cytokines as describe in methods section. The potential association between serum cytokine levels or marker expressing neutrophils frequencies was evaluated by Spearman correlation (one-tailed), with significance defined by a p-value < 0.05: \* for p < 0.05; \*\* for p < 0.01; \*\*\* for p < 0.001; \*\*\*\* for p < 0.0001.

**Figure 2:**
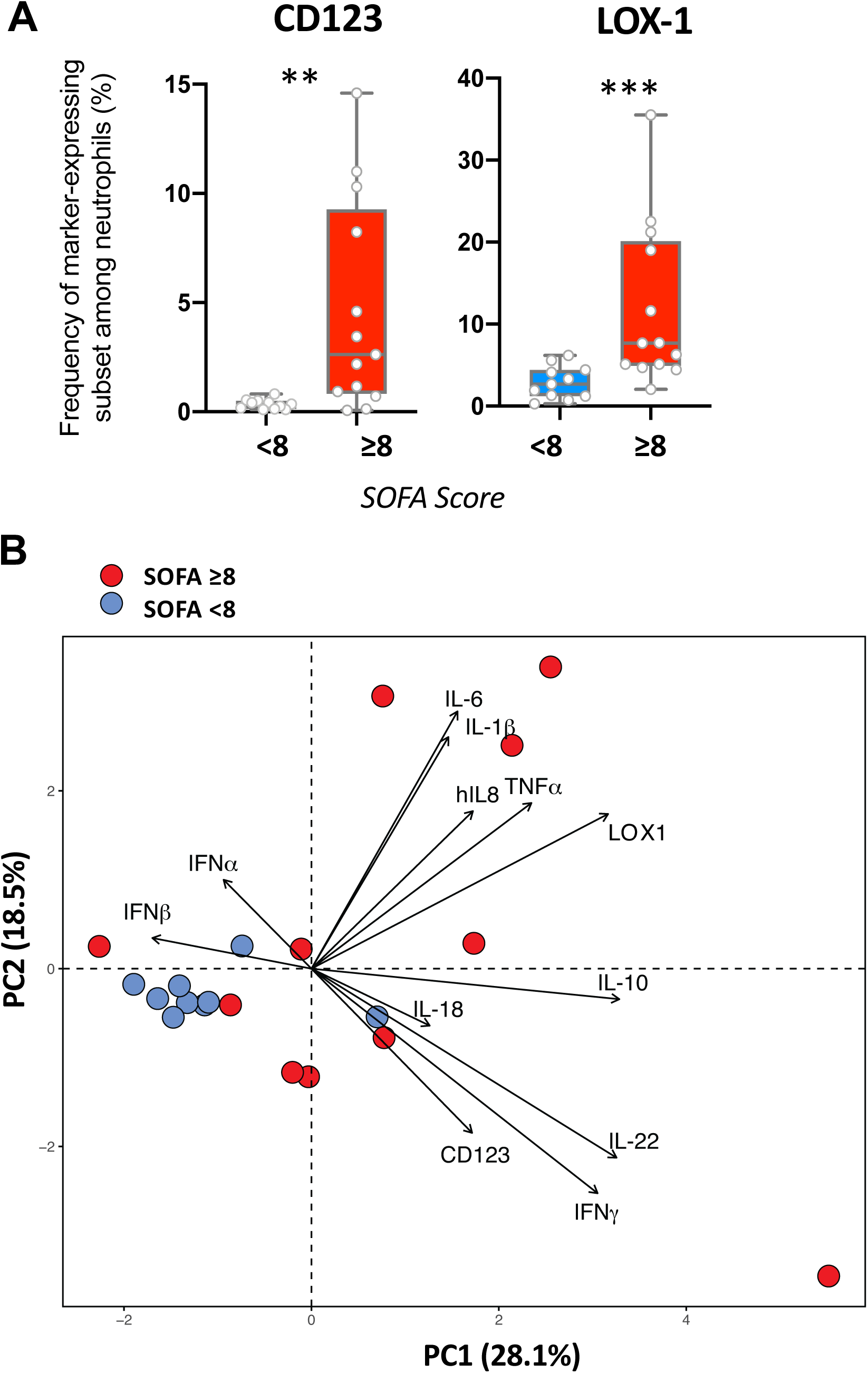
Abundancy of LOX-1-expressing immature neutrophil correlate with clinical severity of COVID-19 patients. (A) Box plots representation (min to max distribution) of the proportion of immature neutrophil expressing CD123- or LOX-1 into patients’ groups with high (n=13) or low SOFA score (n=11) among all ICU patients. Non-parametric two-tailed Mann-Whitney test was used to compare differences in cellular abundance of neutrophil subsets between high and low SOFA score: CD123+ neutrophils (**p<0.0022), LOX-1+ neutrophils (****p<0.0001). (B) Principal component analysis (PCA) using serum cytokines and SOFA score variables on ICU patients sample sizes: high SOFA score (n=11), low SOFA score (n=10) (SOFA<8= blue circles, SOFA≥8 = red circles).

### Increased LOX-1+ immature neutrophil proportions at entry are associated with higher thrombosis risk

Because high values of D-dimer have been associated with disseminated intravascular coagulation in COVID-19 patients (1), we aimed to seek a correlation between the overexpression of LOX-1 and thromboembolic events. The occurrence of thrombosis (Figure 3A) was associated with higher levels of CD123- and LOX-1-expressing immature neutrophils (p<0.01 and p<0.0001 respectively). In order to assess the predictive value of LOX-1 on the occurrence of thrombosis, we performed a Receiver Operating Characteristic (ROC) curve analysis (Figure 3B), with data from patients who survived more that 10 days after inclusion (n=34). The area under the curve (AUC) was 0.95 for LOX-1 immature neutrophil abundancy (p < 0.0001) (Figure 3B) compared with CD123 (AUC=0.81, p=0.003), the SAPS II score (AUC=0.73, p=0.03), D-Dimers level (AUC=0.63, non-significant), and age (AUC=0.61, non-significant). Thus, ROC curve analysis suggested that LOX-1 may be an accurate predictive marker of thrombosis for COVID-19 patients.

**Figure 3:**
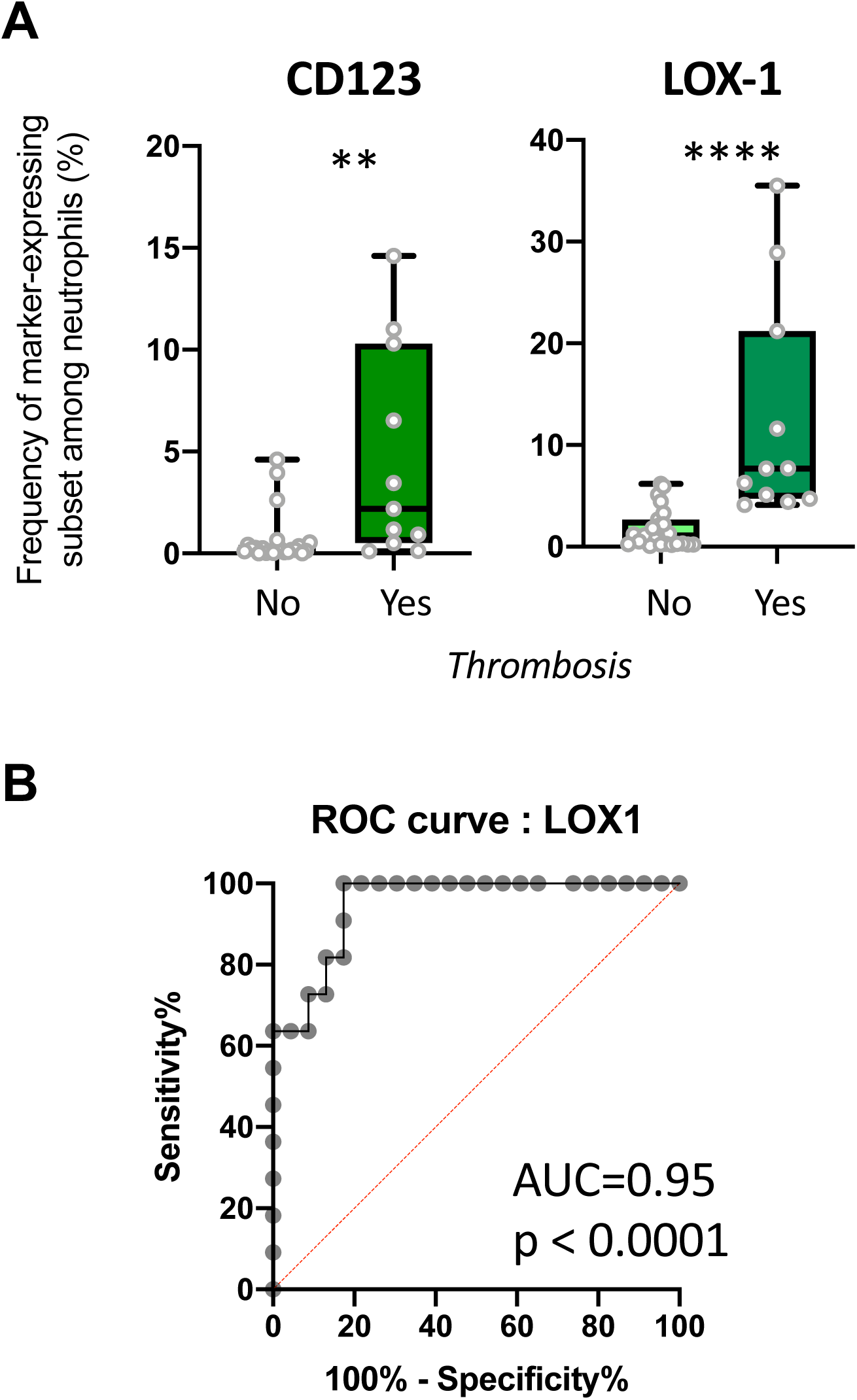
Abundancy of LOX-1-expressing immature neutrophil correlate with thrombosis of critical COVID-19 patients. (A) Box plots representation (min to max distribution) of the proportion of immature neutrophils expressing either CD123- or LOX-1 in blood of patients with (yes) or without (no) thrombosis. Sample sizes: thrombosis n=8, no thrombosis n=24. Nonparametric Mann-Whitney test was used to compare differences in frequencies of neutrophil subsets between the two groups, with significance defined by a p-value < 0.05: * for p < 0.05; ** for p < 0.01; **** for p < 0.0001. (B) Receiver operating characteristic (ROC) curve analysis was performed to assess the predictive value of LOX-1 with thrombosis.

## Discussion

Stratification of patients using biomarkers remains an unmet need in COVID-19 patients care. Here, we sought to identify innate immune cellular signatures that may help predict the outcome of COVID-19 patients with severe symptoms. Conventional whole blood flow cytometry identified classical hallmarks of severe infections, such as neutrophilia and myelemia, but also revealed two novel neutrophils subsets able to discriminate patients requiring ICU or not. Both LOX-1- and CD123-expressing CD10^−^CD64^+^ neutrophil subsets strongly correlated with SAPS II and SOFA severity scores, commonly used in clinical practice for sepsis prognosis. They were also associated with distinct cytokines profiles. Most patients with a high proportion LOX-1 neutrophil subset developed thromboembolic events. The principal limitation of this study is the relatively low number of patients and needs further confirmation in a larger cohort.

Our data indicated that patients’ severity could be predicted based on the proportion of immature CD10^−^CD64^+^ neutrophils using both unsupervised and expert-gating strategies. This neutrophil subset has been described as an immature subset (20) unlike the CD64+-activated neutrophils which still express the neutral endopeptidase (CD10) and the low-affinity immuno-globulin-Fc fragment III (CD16). Numerous studies have shown an association between circulating immature neutrophils and bacterial sepsis (21). Here, we provided evidence that this immature subset may serve as a severity biomarker in COVID-19.

We recently reported that the proportion of CD123-expressing immature neutrophils correlated with bacterial sepsis severity (17). Here, we showed that the expression of CD123 on CD10^−^CD64^+^ neutrophils was related to a higher SOFA score among critically-ill COVID-19 patients. Indeed, both CD123 (the alpha chain of the Interleukin-3 receptor) and its cognate ligand, the IL-3 cytokine, were suggested to play an important role in sepsis. Recent studies demonstrated that a high IL-3 plasma levels were associated with lung inflammation, lung injury and high mortality rates in an animal model, but also in humans (Weber 2015, Tong 2020). In addition, these studies showed that IL3 neutralization and anti-CD123 treatment improved mice outcome by decreasing inflammation, and decreased mortality rates (Weber et al., 2015). IL-3 promotes emergency myelopoiesis, exacerbating pro-inflammatory cytokines secretion and, consequently, systemic inflammation, organ dysfunction and death. The authors further tested the prognostic value of IL-3 in two small cohorts of humans with sepsis and found that high plasma IL-3 levels were associated with high mortality even when adjusting for disease severity. However, we did not observe similar results in COVID-19, as we report an inverse correlation between IL-3 levels and SOFA score. Because IL-3 is mainly produced by activated T-lymphocytes, severe lymphopenia observed in critically-ill COVID-19 patients may limit the IL-3 T cell production.

The association between CD123 expression on immature neutrophils and high serum levels of IL-17, IL-22 and IFNγ was, to the best of our knowledge, never reported and may reveal a yet unidentified link between innate and adaptive immune responses. These findings open the way to new therapeutic opportunities aiming to control the excessive inflammation induced by SARS-CoV-2 infection. The evaluation of CD123 expression on CD10^−^CD64^+^ immature neutrophils could also be a helpful predictor of COVID-19 severity.

The impact of LOX-1 deletion was previously evaluated in a murine model of polymicrobial sepsis, resulting in the reduction of IL-6 and TNFα levels in blood and lungs, enhancing bacterial clearance and preventing neutrophils activation (19). More recently, LOX-1 was identified as a marker on granulocytic myeloid-derived suppressor cells able to suppress T cell activity (18). However, LOX-1 is mostly acknowledged for its role in atherosclerosis. LOX-1 is a class E scavenger receptor contributing to the formation of atherosclerotic plaques by promoting endothelial cell activation, macrophage foam cell formation, and smooth muscle cells migration and proliferation (24). LOX-1 activation induces NFκB activation leading to pro-inflammatory cytokines release, endoplasmic reticulum stress, and reactive oxygen species (ROS) production which could damage the microenvironment (25, 26).

However, LOX-1 role on neutrophils remains elusive. LOX-1 is barely detected on neutrophils at homeostasis, while its expression increases on neutrophils from human cancer patients (18) and in murine sepsis (19, 27).

In this study, LOX-1 expression on immature neutrophils seems to be detrimental for patients as it was associated with the secretion of several pro-inflammatory cytokines, such as IL-6, IL-1β and TNFα, and with severity (as assessed by the SOFA score) and thrombosis. In severe cases of COVID-19, the integrity of the lung is compromised by an exaggerated immune response leading to acute respiratory distress syndrome (10, 16). Mechanisms contributing to microcirculation disorders in sepsis are capillary leakage, leukocytes adhesion and infiltration and intravascular coagulation, leading to thrombus formation. Over the course of systemic inflammatory diseases such as sepsis, the microenvironment is highly oxidative, leading notably to an increase of oxidized low-density lipoprotein (oxLDL) in plasma, which triggers LOX-1 overexpression through a positive feedback loop. In physiological conditions, the increase of LOX-1 expression, especially by endothelial cells, leads to an increase of LDL uptake into vessel wall which activates the specific Oct-1/SIRT-1 thrombosis protective pathway (28). The activation of SIRT1 is able to supress the NFkB-induced expression of tissue factor, also known as thromboplastin, a key initiator of the coagulation cascade involved in thrombus formation (29). In this study, we observed an increase of the incidence of vascular thrombotic events among individuals displaying a high frequency of immature LOX-1^+^ neutrophils. It remains to be seen whether thrombosis in COVID-19 patients results from functionally-diverted neutrophils expressing LOX-1 and/or from its expression on endothelial and smooth muscle cells.

Additionally, we observed a slight correlation between LOX-1 expressing immature neutrophils and the concentration of D-Dimers (spearman test, r=0.42 p= 0.023). D-dimers are used to determine the risk of venous thromboembolism. However, in our study, the predictive score of D-dimers for thromboembolic events was not significant (ROC test, AUC=0.64, p=ns) compared to LOX-1 expressing neutrophil abundancy in the blood (AUC=0.977, p < 0.0001). These results suggest that the high abundancy of LOX-1-expressing immature neutrophils might be sufficient to predict thromboembolic events among critically-ill COVID-19 patients. The overexpression of LOX-1 might also be found in other cell types that might trigger the prothrombotic ERK1/2 pathway. Further investigations would be necessary, such as the titration of oxLDL in blood or the evaluation of the ERK1/2 pathway. In addition, some studies support the relationship between ACE/ACE2 axis and the expression of the pro-oxidative molecule LOX-1, which could increase the oxidative stress favoring prothrombotic state (30). SARS-CoV-2 virus requires binding to ACE2 and is particularly deleterious to patients with underlying cardiovascular disease (31). The polymorphic LOX-1 gene is also intensively associated with increased susceptibility to myocardial diseases. LOX-1 should be thus considered a potential target for therapeutic intervention.

In conclusion, we outline two potential biomarkers of COVID-19 severity measurable among immature neutrophils, the CD123 and LOX-1 surface markers. These markers are significantly correlated with disease severity in general, and more particularly to thromboembolic events. Further research should determine whether an altered neutrophil production is responsible for increased sepsis risk, and how these subsets can be therapeutically targeted.

## Acknowledgments

The authors wish to thank the patients that agreed to participate in this study and all health-care workers involved in the diagnosis and treatment of patients of Assistance Publique Hôpitaux de Paris (AP-HP) and the immunology department staff members. We also thank the administrative staff at the Research and Innovation office (Mrs Laura Wakselman) of Pitié-Salpêtrière Hospital for their support. The study was supported by Fondation de France, « Tous unis contre le virus » framework Alliance (Fondation de France, AP-HP, Institut Pasteur) in collaboration with Agence Nationale de la Recherche (ANR Flash COVID19 program), and by the SARS-CoV-2 Program of the Faculty of Medicine from Sorbonne University (I-COVID programs). LA and PR are recipient of post-doctoral fellowships from the European Union’s Horizon 2020 Research and Innovation Programme under grant agreement No. 681137.

