## Supplementary material for "LOX-1^+^ immature neutrophils predict severe COVID-19 patients at risk of thrombotic complications": table S&

**Supplementary Table S1. Demographics and baseline characteristics of patients with COVID-19**

| | All patients (N=38) | ICU patients (N=24) | Non ICU patients (N=14) |
|---|---|---|---|
| Men | 25 (65.8) | 18 (75) | 7 (50) |
| Age, years, median (range) | 57 (25 – 79) | 55 (25 – 75) | 65 (27 – 79) |
| **Chronic medical illness** | | | |
| Heart disease | 4 (10.5) | 4 (16.7) | 0(0) |
| Type 2 diabetes | 13 (34.2) | 9 (37.5) | 4 (28.6) |
| *Body mass index (kg/m$^2$)* | | | |
| Normal (18.5-25) | 19 (49.7) | 9 (37.5) | 9 (64.3) |
| Overweight (25-30) | 5 (13.2) | 3 (12.5) | 3 (21.4) |
| Obesity (≥30) | 14 (36.8) | 12 (50) | 2 (14.3) |
| Hypertension | 19 (50) | 11 (45.8) | 8 (57.1) |
| Immunocompromised* | 2 (5.3) | 1 (4.2) | 1 (7.1) |
| Malignant tumor | 6 (15.8) | 3 (12.5) | 3 (21.4) |
| Chronic neurologic disease | 1 (2.6) | 1 (4.2) | 0 (0) |
| Chronic pulmonary disease | 5 (13.2) | 4 (16.7) | 1 (7.1) |
| Chronic kidney disease | 6 (15.8) | 3 (12.5) | 3 (21.4) |
| Chronic liver disease | 0 (0) | 0 (0) | 0 (0) |
| *Smoking habits* | | | |
| Never smoker | 31 (81.6) | 21 (87.5) | 10 (71.4) |
| Former smoker | 4 (10.5) | 3 (12.5) | 1 (7.1) |
| Daily smoker | 3 (7.9) | 0 (0) | 3 (21.4) |
| *Past history of arterial or venous thrombosis* | 6 (15.8) | 4 (16.7) | 2 (14.3) |
| Arterial | 5 (13.2) | 4 (16.7) | 1 (7.1) |
| Venous | 1 (2.6) | 0 (0) | 1 (7.1) |
| **Treatment regimen at baseline** | | | |
| Long-term immunosuppressive agent use | 1 (2.6) | 1 (4.1) | 0 (0) |
| Glucocorticoids | 1 (2.6) | 1 (4.2) | 0 (0) |
| Nonsteroidal anti-inflammatory drugs | 0 (0) | 0 (0) | 0 (0) |
| Recent chemotherapy for cancer | 2 (5.3) | 1 (4.2) | 1 (7.1) |
| Angiotensin converting enzyme inhibitor | 10 (26.3) | 6 (25) | 4 (28.6) |
| Angiotensin II receptor blockers | 6 (15.8) | 4 (41.7) | 4 (12.3) |
| Oral anticoagulant | 0 (0) | 0 (0) | 0 (0) |
| **Severity score at baseline** | | | |
| SAPS II, median (range) | 33 (25 - 78) | 35.5 (15 – 78) | 25.5 (9 – 61) |
| SOFA, median (range) | | 8.5 (2 – 17) | |
| **Time from onset of symptoms to admission** | | | |
| Days, median (range) | 8 (5 – 47) | 8 (5 – 22) | 13 (1 – 47) |
| **Laboratory findings at baseline** | | | |
| Leucocytes, x10$^9$/L, median (range) [normal range: 4.0-10.0] | 9.29 (1.19 – 23.79) | 10.3 (3.43 – 23.79) | 7.705 (1.35 – 19.86) |
| Neutrophil count, x10$^9$/L, median (range) [normal range : 2.7 – 5] | 7.87 (1.35 – 60.76) | 8.75 (2.72 – 60.76) | 7.705 (1.35 – 19.86) |
| Lymphocyte count, x10$^9$/L, median (quartiles) [normal range: 1.5 – 4] | 0.94 (0.56 – 1.34) | 0.925 (0.48 – 2) | 1.185 (0.24 – 1.81) |
| Lactate dehydrogenase, U/L, median (range) [normal range: 135-215] | 475.5 (234 – 2030) | 504 (375 – 1087) | 324 (234 – 2030) |
| Ddimer, ng/mL, median (range) | 2450 (540 – 20000) | 2760 (540 – 20000) | 1860 (540 – 20000) |

| | | | |
|---|---|---|---|
| **Chest CT finding: extension of GGO and/or consolidation[¤]** | | | |
| 0% | 0 (0) | 0 (0) | 0 (0) |
| <10% | 4 (14.3) | 0 (0) | 4 (33.3) |
| 10-25% | 4 (14.3) | 1 (6.3) | 3 (25) |
| 25-50% | 6 (21.4) | 2 (12.5) | 4 (33.3) |
| 50-75% | 9 (32.1) | 8 (50) | 1 (8.3) |
| > 75% | 5 (17.9) | 5 (31.3) | 0 (0) |
| **Treatment** | | | |
| *Hydroxychloroquine* | 16 (42.1) | 14 (58.3) | 2 (14.3) |
| *Glucocorticoids* | 1 (2.6) | 1 (4.2) | 0 (0) |
| *Tocilizumab or sarilumab* | 0 (0) | 0 (0) | 0 (0) |
| *Oseltamivir* | 5 (13.2) | 5 (20.8) | 0 (0) |
| *Antibiotic therapy* | 38 (100) | 24 (100) | 14 (100) |
| *Oxygen therapy* | 38 (100) | 24 (100) | 14 (100) |
| Nasal cannula | 14 (36.8) | 0 (0) | 14 (100) |
| Non-invasive ventilation or high-flow nasal cannula | 3 (7.9) | 3 | 0 |
| Invasive mechanical ventilation | 21 (55.3) | 21 (87.5) | 0 (0) |
| *Extracorporeal membrane oxygenation* | 13 (34.2) | 13 (54.2) | 0 (0) |
| *Hemodialysis* | 10 (26.3) | 8 (33.3) | 2 (14.3) |
| **Complications** | | | |
| Acute respiratory distress syndrome | 21 (55.3) | 21 (87.5) | 0 (0) |
| Acute kidney injury | 12 (31.2) | 11 (45.8) | 1 (7.1) |
| Pulmonary embolism | 2 (5.3) | 1 (4.2) | 1 (7.1) |
| Thrombosis | 10 (26.3) | 10 (41.7) | 0 (0) |
| Venous | 10 (26.3) | 10 (41.7) | 0 (0) |
| Arterial | 0 (0) | 0 (0) | 0 (0) |
| **Clinical outcome[†]** | | | |
| Discharged | 29 (76.3) | 13 (54.2) | 14 (100) |
| Remained in hospital | 4 (10.5) | 4 (16.7) | 0 (0) |
| Death | 5 (13.2) | 5 (20.8) | 0 (0) |

Values are expressed as n (%), unless stated otherwise.
*including cardiac, liver or kidney allograft, hematopoietic stem cell transplantation, or immunosuppressive agent for auto-immune disease
[¤] 28 patients were assessed
[†] As of June 8th, 2020
CT, computed tomography; GGO, ground-glass opacities; SAPS II, Simplified Acute Physiology Score II; SOFA score, Sequential organ failure assessment score

**Supplementary table S2**

| REAGENT or RESOURCE | SOURCE | IDENTIFIER |
|---|---|---|
| **Antibodies** | | |
| CRTH2 FITC | Biolegend | #350107 |
| CD123 PE | Biolegend | #306005 |
| LOX1 BV421 | Biolegend | #358609 |
| CD64 BV605 | Biolegend | #305033 |
| PDL1 BV711 | Biolegend | #329721 |
| CD15 BV786 | BD | #741013 |
| CD14 BUV737 | BD | #564444 |
| CD10 BUV395 | BD | #565975 |
| Brillant violet buffer | BD | #563794 |
| **Critical Commercial Assays** | | |
| Human CorPlex™ Cytokine Panel 7-Plex array | Quanterix | # 85-0410 |
| Simoa™ Human IL-3 Discovery Kit | Quanterix | #102462 |
| Simoa™ IL-17A Advantage Kit | Quanterix | #101599 |
| Simoa™ IL-18 Discovery Kit | Quanterix | #102700 |
| Simoa™ GM-CSF Advantage Kit | Quanterix | #102329 |
| Simoa™ Human IFN-a Advantage Kit | Quanterix | #100860 |
| VeriKine-HS™ Human IFN Beta ELISA Kit | PBL Assay Science | #41415 |
| **Software and Algorithms** | | |
| RStudio version 1.3.959 Mac | Opensource | https://rstudio.com/ |
| Cytobank | Cytobank | https://inserm.cytobank.o |
| Prism version 8.00 | Graphpad | https://www.graphpad.co |
| SP-X Analysis Software | Quanterix | https://www.quanterix.co |
| Simoa HD-1 Software | BD | https://www.flowjo.com/ |
