## Supplementary material for "LOX-1^+^ immature neutrophils predict severe COVID-19 patients at risk of thrombotic complications": Figures S1 and S2

<u>**Supplementary Figures**</u>

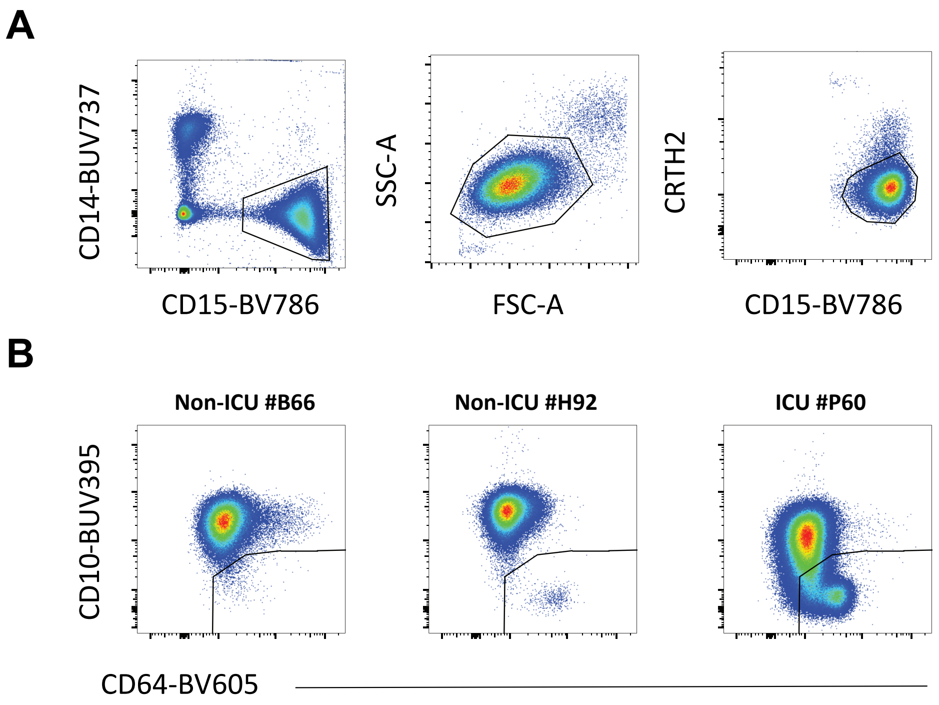


**Supplemental Figure S1: Gating Strategy** used for the analysis of neutrophil populations. After debris and doublets exclusion, $CD15^{+}CD14^{-}$ cells were selected. Eosinophils were excluded through SSC/FSC parameters and the expression of CRTH-2. The proportion of immature neutrophils was evaluated through the expression of CD10 and CD64. One representative individual of uninfected donor (#B66), COVID-19 non IUC patient (#H92) and COVID-19 ICU patient (#60) are shown as an example.

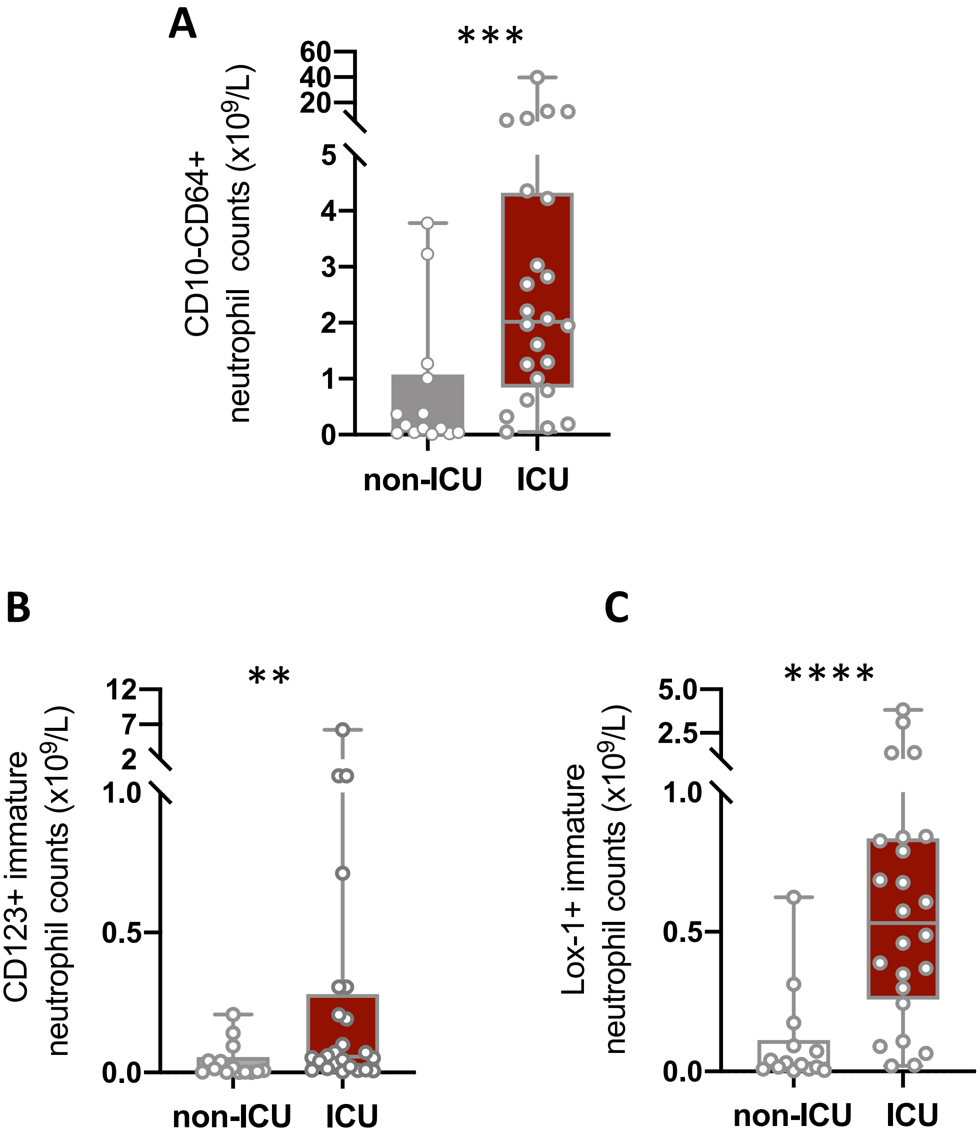


**Supplementary Figure S2: Critical COVID-19 patients displayed increased numbers of immature neutrophil subsets expressing CD123 or LOX-1.** Box plots representation (min to max distribution) of A) CD10$^-$CD64$^+$ neutrophil B) CD123+ immature CD10-CD64+ neutrophil and C) LOX-1 immature CD10-CD64+ neutrophil cell counts in ICU and non-ICU patients' groups. identify the median and min to max distribution. Nonparametric Mann-Whitney test was used to compare differences in neutrophil subset cell counts between groups, with significance defined by a p-value < 0.05: * for p < 0.05; ** for p < 0.01; *** for p < 0.001.
