## Supplementary material for "LOX-1^+^ immature neutrophils predict severe COVID-19 patients at risk of thrombotic complications": sup methods

**Supplementatl methods**

**Quanterix technology (digital ELISA)**

The Simoa$^{TM}$ (single molecule array) HD-1 analyser (Quanterix, Lexington, MA, USA) was used for ultrasensitive immunodetection (digital ELISA) of IL-3, IL-17A, IL-18, GM-CSF and IFN-α, using singleplex bead-based assays (Supplemental Table S2). Concentrations of IL-1β, IFN-γ, IL-6, IL-8, IL-22, TNF-α and IL-10 were determined using a multiplex planar array immunoassay on the Quanterix SP-X™ platform according to manufacturer's instructions. Serum IFN-β levels were quantified with a highly sensitive ELISA kit (PBL Assay Science, Piscataway, NJ, USA). The concentrations of cytokines in unknown samples were interpolated from a standard curve performed with two replicates of each level of recombinant calibrator proteins, representing the dynamic range of the assay: IL-1β (0.073-300 pg/mL), IFN-γ (0.012-50 pg/mL), IL-6 (0.073-300 pg/mL), IL-8 (0.098-400 pg/mL), IL-22 (0.024-100 pg/mL), TNFα (0.098-400 pg/mL), IL-10 (0.024-100 pg/mL), IL-3 (0.686-500 pg/mL), IL-17A (0.041-30 pg/mL), IL-18 (0.011-45 pg/mL), GM-CSF (0.041-30 pg/mL), IFN-α (0.028-27.3 pg/mL) and IFN-β (1.2-150 pg/mL).

**Computational data analysis of neutrophils**

viSNE analysis was performed with Cytobank software (Supplemental Table S2) using CD10, CD64, PD-L1, CD123 and LOX-1 markers. We selected a total of 100,000 CD15+ granulocytes (following the gating strategy in Supplementary Figure 1), divided equally between 24 ICU patients and 13 non-ICU patients, using one single concatenated file for each condition.
